# Conformational Dynamics of Na^+^-Pumping NADH-Quinone Oxidoreductase during Na^+^ Translocation from AlphaFold-Facilitated Markov State Modeling

**DOI:** 10.64898/2026.01.28.702413

**Authors:** Takehito Seki, Jun Ohnuki, Kei-ichi Okazaki

## Abstract

The Na^+^-pumping NADH–quinone oxidoreductase (Na^+^-NQR) is a respiratory chain enzyme found in pathogenic bacteria, including *Vibrio cholerae*, and is essential for energy metabolism by generating a transmembrane Na^+^ gradient that drives ATP synthesis and flagellar motility. Because the molecular structure of Na^+^-NQR is unrelated to the corresponding mitochondrial H⁺-pumping NADH-quinone oxidoreductase (respiratory complex I), it is a promising antibiotic target. Although it has been shown that Na^+^ pumping is mediated by an alternating-access conformational change in the NqrD/E subunits, coupled to redox switching of a cofactor, the thermodynamics and kinetics of the conformational transition, including the free-energy profile and the rate-limiting steps, remain unclear. Here, we construct redox-state-dependent Markov state models (MSMs) from extensive molecular dynamics (MD) trajectories in the oxidized and reduced states to quantify the conformational free-energy landscapes and primary transition pathway. To accelerate conformational sampling, MD simulations are initiated from diverse NqrD/E conformations generated by AlphaFold. Our analysis clarifies how the NqrD/E conformation is regulated by the redox state and by Na^+^ binding to achieve Na^+^ translocation. This study provides a quantitative framework for understanding ion-pumping mechanisms of redox-driven membrane proteins.

## Introduction

The Na^+^-pumping NADH-quinone oxidoreductase (Na^+^-NQR) is a respiratory chain enzyme found in pathogenic bacteria, including *Vibrio cholerae*. Na^+^-NQR catalyzes electron transfer from nicotinamide adenine dinucleotide (NADH) to quinone, coupled to Na^+^ efflux from the cell, generating a Na^+^ gradient across the cellular membrane. This gradient is essential for the microbial energy-requiring processes such as ATP synthesis, flagellar motility, and nutrient uptake.^1–3^ In mammalian cells, the corresponding respiratory chain enzyme is known as mitochondrial H⁺-pumping NADH-quinone oxidoreductase (respiratory complex I). These respiratory chain enzymes share the functional property of using electron transfer to drive cation pumping. However, they are structurally and evolutionarily unrelated, which makes Na⁺-NQR a promising target for highly selective antibiotics.^3,4^

X-ray crystallography^5,6^ and cryogenic electron microscopy (cryo-EM)^6,7^ have revealed that the entire architecture of Na^+^-NQR consists of six subunits (NqrA - NqrF) and cofactors (riboflavin derivatives and 2Fe-2S clusters) embedded within the subunits (Figure 1A). The NqrF subunit contains a Rossmann nucleotide binding motif to which NADH binds transiently, from which electron is initially transferred to the non-covalently bound flavin adenine dinucleotide: FAD (FAD^NqrF^). The NqrF subunit also contains a 2Fe–2S cluster (2Fe-2S^NqrF^), which is the next acceptor of electrons. Multiple conformations of the NqrF subunit have been observed by cryo-EM (PDB IDs: 9UD6, 9UD8, and 9UD9).^8^ Particularly in the conformation closest to the membrane-embedded NqrD/E subunits (PDB ID: 9UD9), which contain another 2Fe-2S cluster (2Fe-2S^NqrD/E^), the distance between 2Fe-2S^NqrF^ and 2Fe-2S^NqrD/E^ is relatively short (∼16 Å; ∼30 Å in most experimental conformations), suggesting an inter-subunit electron transfer pathway from 2Fe-2S^NqrF^ to 2Fe-2S^NqrD/E^. Subsequent inter-subunit electron transfer to a flavin mononucleotide: FMN (FMN^NqrC^) requires conformational rearrangements of the NqrC subunit, observed in some structures (PDB IDs: 8ACW and 8ACY). Electrons are further transferred through FMN^NqrB^ and Riboflavin: RBF (RBF^NqrB^) in the NqrB subunit, with RBF^NqrB^ being close to the quinone binding site, where electrons are finally passed to ubiquinone. The NqrA-NqrB interface is involved in quinone binding: cryo-EM confirmed that quinone derivatives exhibit an inhibitory activity of Na^+^-NQR by binding to the interface.^9–12^ The cofactor-mediated electron transfer (NADH → FAD^NqrF^ → 2Fe-2S^NqrF^ → 2Fe-2S^NqrD/E^ → FMN^NqrC^ → FMN^NqrB^ → RBF^NqrB^ → ubiquinone) is coupled to Na^+^ translocation, which has been observed in biochemical and biophysical studies.^7,13^ However, the transmembrane subunits involved in the Na⁺ translocation had been controversial.^5,6,13,14^

**Figure 1.**
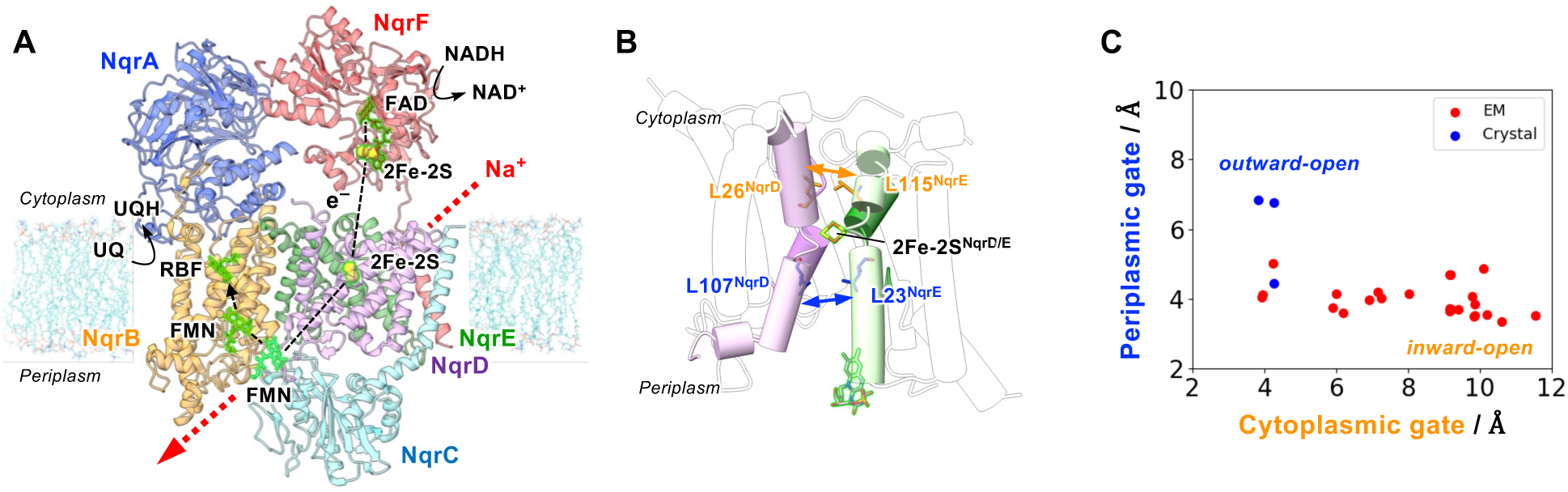
Structural information of Na^+^-NQR. (A) Overall structure of Na^+^-NQR shown in cartoon representation (PDB ID: 7XK3). Electron transfer via cofactors is represented by the purple broken line. (B) The transmembrane helices in NqrD/E are shown with the hydrophobic gate residues (inward gate: Leu26^NqrD^-Leu115^NqrE^ and outward gate: Leu107^NqrD^-Leu23^NqrE^) that control access to the Na^+^ binding site from either the inward (cytoplasmic) or outward (periplasmic) side of the membrane. The inward and outward gates are represented in orange and blue sticks, respectively. (C) The cytoplasmic and periplasmic gate sizes in experimental structures listed in Figure S2.

Recently, we combined cryo-EM and all-atom molecular dynamics (MD) simulations to investigate how electron transfer within Na⁺-NQR induces structural rearrangements in protein subunits and drives ion translocation.^8^ MD simulations provided atomistic insight into how the protein fulfills its function through conformational dynamics. The MD simulations in which the redox states of the cofactors were switched showed that the reduction of 2Fe-2S^NqrD/E^ induces Na⁺ binding to a nearby transmembrane site. Subsequently, the targeted MD simulations demonstrated that an alternating-access conformational change in the NqrD/E subunits enables Na⁺ translocation, as in other transporter proteins.^15–17^ However, we have not identified how the alternating-access conformational change underlying Na⁺ translocation is triggered, the intermediate states during translocation, including the rate-limiting steps, and the free energy profile of the conformational change.

Markov state modeling (MSM)^18–21^ enables quantitative characterization of the thermodynamics and kinetics of underlying dynamic processes. MSM can also provide transition pathways, including key transition states that determine reaction rates and mechanisms. Because MSM construction requires a large number of computationally expensive MD trajectories, efficiently generating them remains a central challenge. To address this, we have leveraged the AI-based AlphaFold (AF)^22,23^ to generate diverse initial protein conformations for MD simulations.^24^ AF extracts coevolutionary coupling between amino acid positions from multiple sequence alignment (MSA) of a query sequence, thereby inferring spatial proximity of residue pairs in native protein structures. MSA-related methods such as MSA subsampling,^25^ clustering,^26^ and masking,^27^ which restrain the amount of coevolutionary coupling information during structure inference, have successfully generated not only stable but also metastable or intermediate conformations. Using such conformations as initial structures for MD simulations accelerates conformational sampling.^24,28^

In this study, we construct redox-state-dependent MSMs using extensive MD trajectories initiated from AF-predicted diverse protein conformations to elucidate conformational free energy and pathway of Na⁺-NQR during ion translocation coupled to electron transfer. In the previous work, we identified that Na⁺-NQR mediates ion translocation across the membrane through an alternating-access conformational change in the NqrD/E subunits.^8^ To elucidate the detailed mechanism, we first generate a diverse set of NqrD/E conformations with modified AF3 protocols.^29^ We then perform MD simulations in two different redox states initiated from both experimental structures and AF-generated models, followed by adaptive sampling cycles in underexplored conformational space.^30^ The resulting redox-state-dependent MSMs identify the thermodynamics, driving forces, and primary transition pathways associated with the conformational changes in the NqrD/E subunits underlying Na⁺ translocation. We show that the free energy of the alternating-access conformational change depends on both the redox state of 2Fe-2S^NqrD/E^ and the Na⁺-binding state, demonstrating that Na⁺-NQR achieves the ion translocation by switching these chemical states.

## Results

### Conformational sampling by AlphaFold3

First, we generated a wide-range of Na^+^-NQR conformations using AlphaFold3 (AF3). All six subunits (NqrA - NqrF) were included with six cofactor molecules for the AF3 prediction. According to our previous work,^8^ we focus on the NqrD and NqrE subunits as the main components that mediate Na^+^ translocation through an alternating-access conformational change (Figures 1B, 1C, and S2). The distances of the cytoplasmic gate (Leu26^NqrD^ - Leu115^NqrE^) and the periplasmic gate (Leu107^NqrD^ - Leu23^NqrE^) were monitored to assign the conformational state. The MSA subsampling approach^25^ was initially adopted to enhance conformational sampling, where the MSA depth, fixed in the original AF3 code, can be specified as an option in our implementation (af3_mmm).^29^ With the default MSA depth, the predicted structures were concentrated in a slightly outward-open conformation (Figure 2A). In contrast, predictions with shallower MSA depths generated wider outward-open conformations (Figure 2B). Because the MSA subsampling was unable to generate the other conformational states in this case, we attempted AF3 prediction using template structures. Here, using experimentally determined inward-open conformations (PDB IDs: 7XK3 and 9UD9) as templates, we generated conformations spanning the occluded-to-inward-open states (Figure 2C). In this way, we have successfully covered diverse Na^+^-NQR conformations leveraging AF3, which will serve as the initial structures for MD simulations.

**Figure 2.**
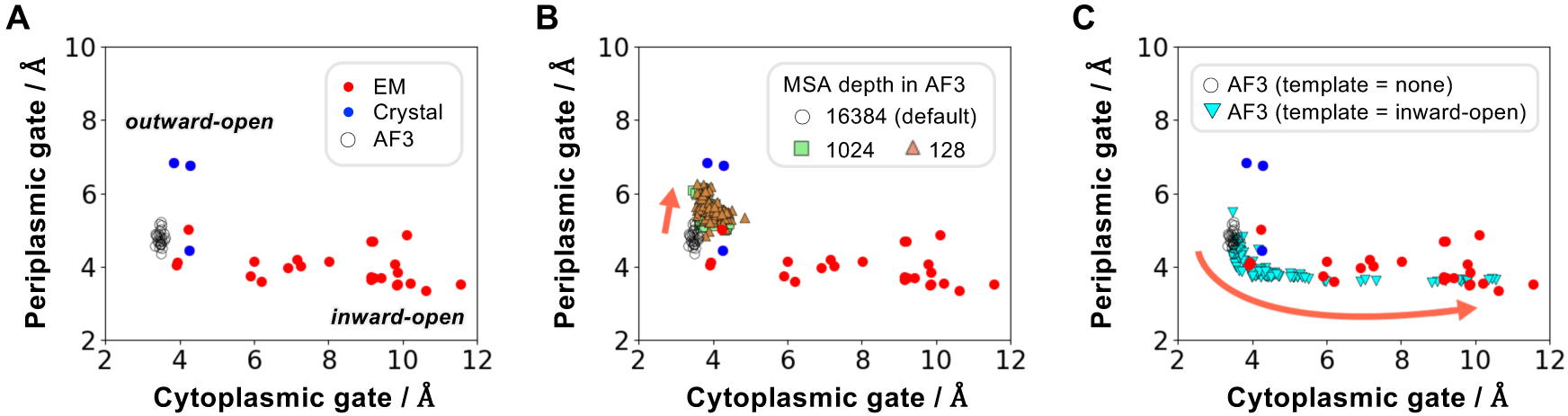
Conformational sampling of Na^+^-NQR by AlphaFold3. (A-C) Structural data plotted on the axis defined by the cytoplasmic and periplasmic gate sizes in NqrD/E. AlphaFold3 with the default MSA depth (A), with shallower MSA depths (B), and with an inward-open conformation as the template (C) generated 25, 100, 100 structures, respectively.

### AlphaFold-facilitated Markov state modeling with adaptive sampling

To elucidate the thermodynamics, driving forces, and primary pathways involved in the alternating-access conformational change, we develop an AlphaFold-facilitated Markov state modeling (MSM) framework based on MD trajectories initiated from AF predictions described above. The redox-state-dependent MSMs are constructed from MD simulations in specific redox states, as described in subsequent sections. In addition to AF and experimental structures, we initiate MD trajectories based on an adaptive sampling approach.^30^ The overall workflow is illustrated in Figure 3. The MD trajectories initiated from the experimental and AF structures for reduced 2Fe-2S^NqrD/E^ were listed in Table S1. Each MD trajectory was then described using input features, which characterize the conformational changes in NqrD/E. The input features were determined via contact analysis as explained in the Methods and Table S2. These features were then subjected to time-lagged independent component analysis (tICA) for dimensionality reduction, and the resulting projections were clustered using the k-means algorithm to define discrete microstates. In the adaptive sampling procedure, additional MD trajectories were iteratively generated by restarting MD simulations from structures in the least-visited microstates, thereby improving sampling efficiency of underexplored regions in conformational space. The three cycles of adaptive sampling yielded a total of 7.8 µs MD trajectories for the reduced 2Fe-2S^NqrD/E^ (Table S1 and Table S3), which were used to construct a final MSM. In the final MSM, the MD trajectories were discretized into 100 microstates with the first 20 tICA modes (tICs). The numbers of microstates and tICs were determined according to the VAMP-2 score^31^ (Figure S1B). The implied timescale converged at an MSM-lag time of 4 ns, which was therefore used for the subsequent MSM analysis (Figure S1C). The free energy was calculated from the equilibrium probabilities of the first MSM mode with the eigenvalue *λ_i_* = 1 (Figure S1D).

**Figure 3.**
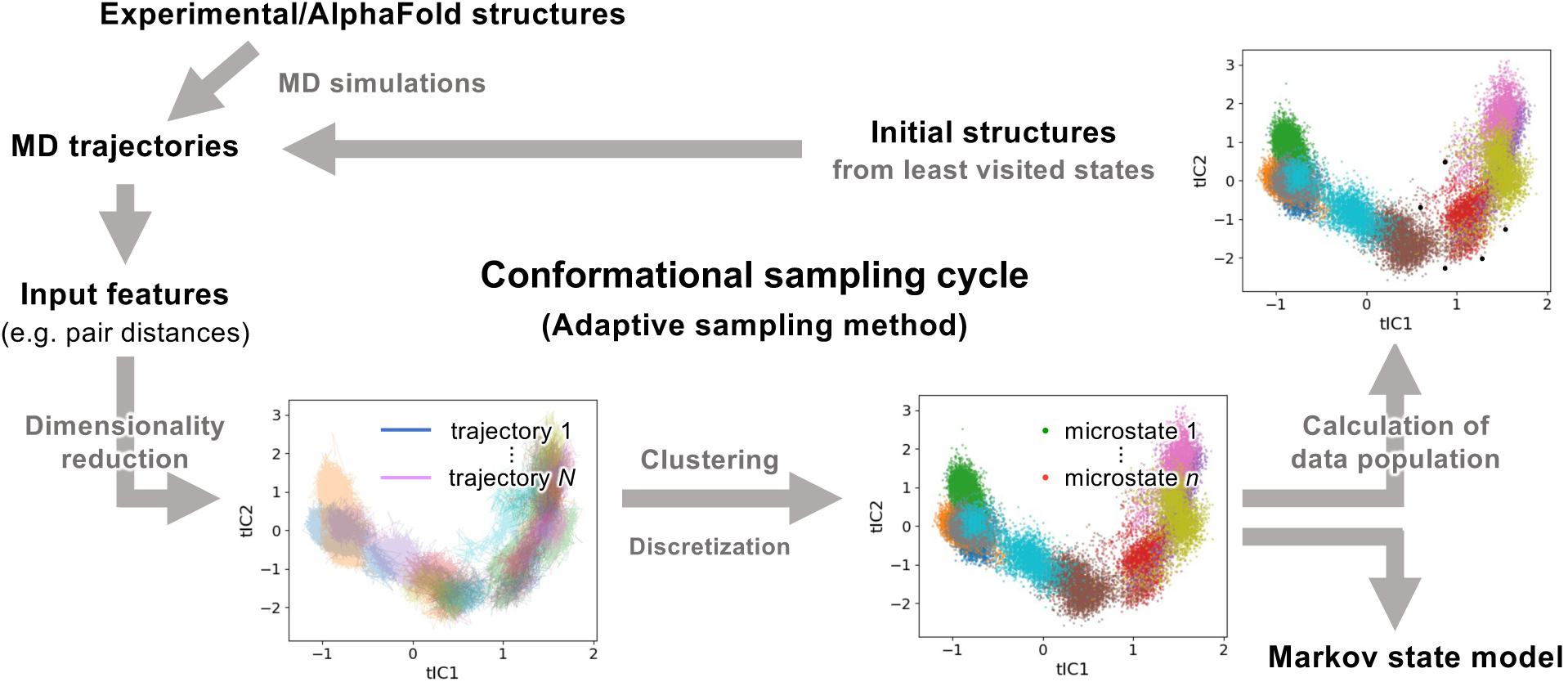
Workflow for conformational sampling via the adaptive sampling approach.

### Markov state modeling of the reduced 2Fe-2S^NqrD/E^ state

Using MD trajectories under the reduced 2Fe-2S^NqrD/E^, we constructed an MSM and evaluated the conformational free energy of NqrD/E with the inward (cytoplasmic) and outward (periplasmic) gate lengths as the axes (Figure 4A). The free energy landscape indicates that the inward-open conformation is the most stable state, whereas the occluded and outward-open conformations are metastable. When the free energy landscape was plotted with the cytoplasmic gate length and z-coordinate of the Na^+^ being translocated in the NqrD/E transmembrane region, it showed that Na^+^ binding is stabilized near 2Fe-2S^NqrD/E^ and is coupled to closing of the cytoplasmic gate (Figure 4B). This suggests that the Na^+^ binding contributes to the conformational change in NqrD/E from the inward-open to the occluded state. To obtain a clear understanding, the free energy landscapes were plotted separately for the Na^+^-unbound and Na^+^-bound states, as shown in Figures 4C and 4D, respectively. These plots clarified that the occluded state (defined as cytoplasmic gate < 5 Å and periplasmic gate < 5 Å) is unstable in the Na^+^-unbound state, whereas it becomes stable with Na^+^ bound. This suggests that a negative charge of the reduced 2Fe-2S^NqrD/E^ must be neutralized by Na^+^ in the hydrophobic transmembrane region to drive the conformational changes from the inward-open to the occluded/outward-open states.

**Figure 4.**
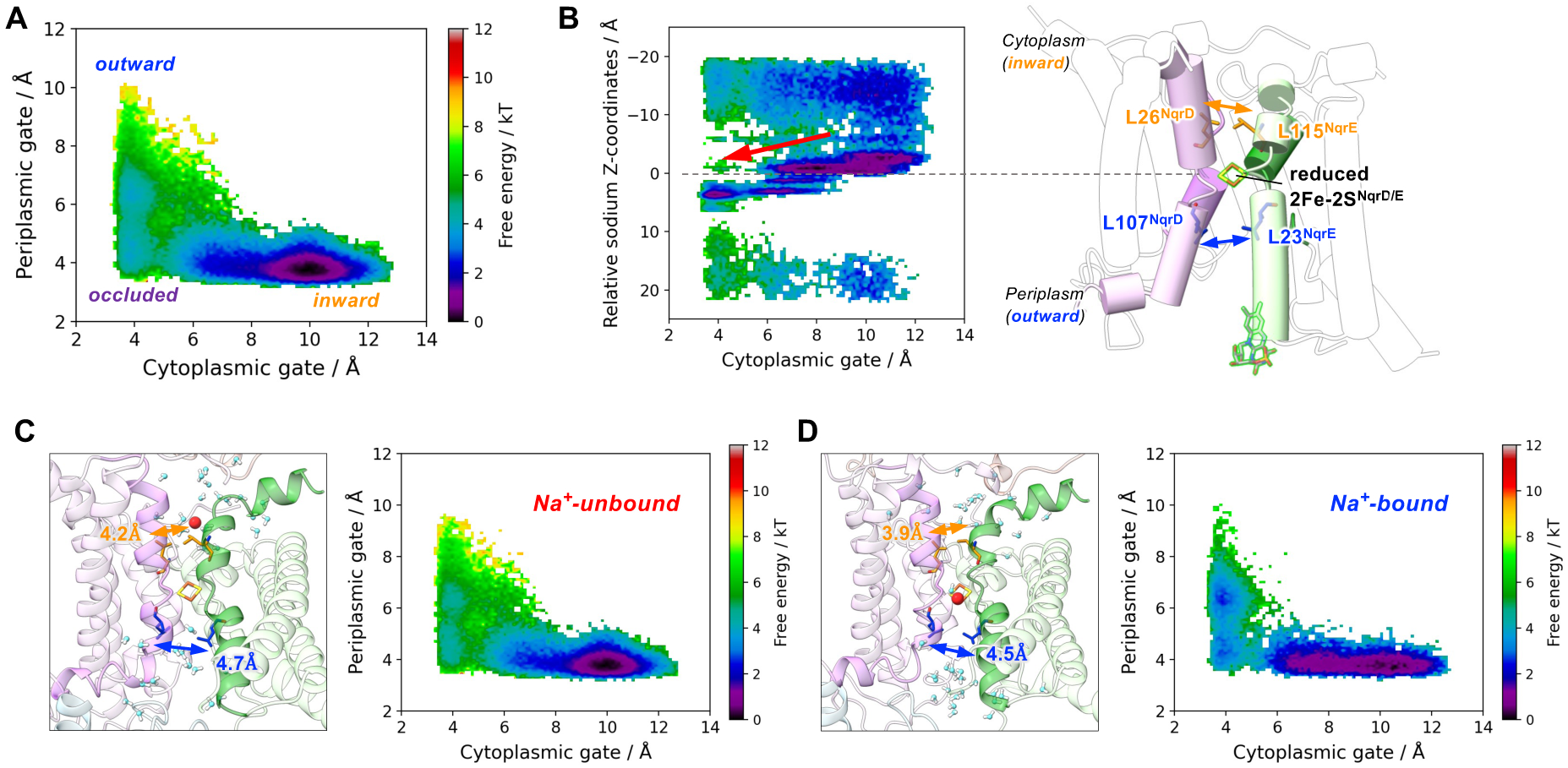
Free-energy analysis of NqrD/E conformational changes under the reduced 2Fe-2S^NqrD/E^ condition. (A) Conformational free energy landscape projected on the axes of the cytoplasmic and periplasmic gate sizes. (B) Projection on the cytoplasmic gate size and relative z-coordinates of the Na^+^ being translocated in the transmembrane NqrD/E (*z_Na_+*). (C-D) Representative snapshots of the occluded state and free energy landscapes with the gate axes under the Na^+^-unbound condition (|*z_Na_+*| > 8 Å) (C), or under the Na^+^-bound condition (|**z*_Na_+*| < 8 Å) (D). Na^+^ ions and water molecules are represented as red spheres and cyan/white ball-and-sticks, respectively.

### Transition path analysis of the reduced 2Fe-2S^NqrD/E^ state

To analyze the transition pathways, the 100 discrete microstates defined in the MSM were coarse-grained into 31 macrostates using the PCCA+ algorithm,^32^ and transition path analysis was performed using these macrostates. Here, the inward-open conformations were defined as the initial state and the outward-open conformations as the final state, and the reactive fluxes between states were computed.^33,34^ As a result, the primary transition pathway accounts for 14 % of the total flux (see Methods). Representative structural snapshots, distributions of the Na⁺ z-coordinate in the transmembrane region, and the committor values along the primary transition pathway are shown in Figure 5. The committor value, ranging from 0 to 1, represents the progress of the conformational change from the inward-open to the outward-open state.^33–36^ Here, the analysis showed that the conformational changes in NqrD/E are coupled to the translocation of intracellular Na^+^ across the membrane, consistent with the previous targeted MD result.^8^ From the initial state (state ID 0) to state IDs 29 and 26, the committor value remained nearly zero, whereas intracellular Na^+^ influx into the transmembrane region was observed. When the committor value increased slightly (to 0.07), a conformational transition to the occluded state was observed with the reduced cytoplasmic gate size to ∼5 Å, accompanied by a shift in the Na^+^-binding site toward the extracellular side with the z coordinate of Na^+^ less than zero. These observations suggest that Na^+^-binding drives the NqrD/E conformational changes from the inward-open to the occluded state, consistent with the free-energy analysis in the previous section. As the committor value increased further, the conformational transition to the outward-open conformations and Na^+^ efflux to the extracellular side were observed. During this transition, a transient approach of 2Fe–2S^NqrD/E^ toward FMN^NqrC^ was also observed (Figure S3), consistent with our previous study.^8^ In the final outward-open state (state ID 30), the presence of Na^+^ on the extracellular side was observed again, suggesting transient rebinding of Na⁺. The smallest reactive fluxes along the primary pathway were those from the occluded state to the outward-open state (IDs 21 to 11 or 11 to 4), indicating that these steps are rate-limiting. The rate-limiting steps involve the opening of the periplasmic gate (Figure S3). Furthermore, the mean first-passage time (MFPT) from the inward-open (ID 0) to the outward-open (ID 30) states was approximately 1.5 ms (376,168 MSM steps × 4 ns MSM lag time).

**Figure 5.**
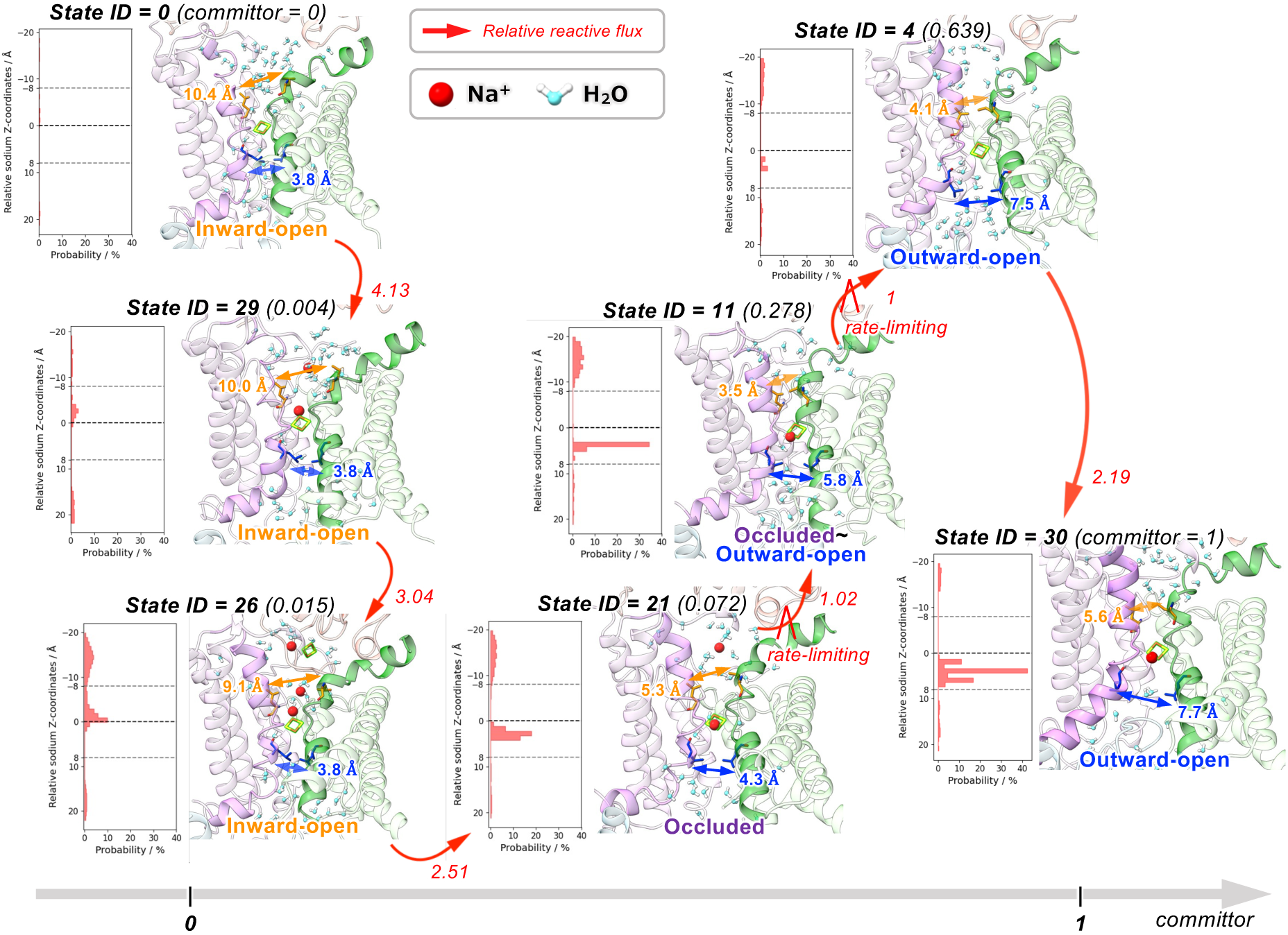
The primary transition pathway from the inward-open to the outward-open state. The relative dominance of the transition pathways was estimated from the reactive fluxes among 31 coarse-grained states. Each net-reactive flux is normalized by the minimum (rate-limiting) net-reactive flux (from state 11 to 4), and shown in red next to an arrow. For each state, a representative structural snapshot, distribution of the Na⁺ z-coordinate in the transmembrane region, and the committor value are shown.

### Markov state modeling of the oxidized 2Fe-2S^NqrD/E^ state

The MSM of the reduced 2Fe-2S^NqrD/E^ suggested that the Na^+^-binding near the 2Fe-2S^NqrD/E^ drives the conformational changes in NqrD/E from the inward-open to the occluded/outward-open states. Here, we investigated the effect of 2Fe-2S^NqrD/E^ redox state on conformational free energy of NqrD/E by constructing an MSM of the oxidized 2Fe-2S^NqrD/E^. An MSM was constructed from MD trajectories under the oxidized 2Fe-2S^NqrD/E^ condition. The representative structures from each of the 30 macrostates identified in the previous MSM (reduced 2Fe-2S^NqrD/E^) were used as the initial structures for MD simulations with the redox state of 2Fe-2S^NqrD/E^ switched to the oxidized state. Based on the resulting MD trajectories (40 ns × 30 = 1.2 µs), the adaptive sampling approach was applied to generate a total of 8.2 µs of trajectories across three sampling cycles (Figure 3). The free energy landscapes calculated from the MSM are shown in Figure 6. In the oxidized state, stable Na^+^-binding was not observed near the 2Fe-2S^NqrD/E^ (Figure 6A), consistent with our previous work.^8^ Projecting on the cytoplasmic and periplasmic gate sizes, it was revealed that the occluded state is also thermodynamically stable in addition to the inward-open state (Figure 6B). Unlike the reduced form, the oxidized 2Fe-2S^NqrD/E^ has no negative charge and therefore does not destabilize the hydrophobic transmembrane region of NqrD/E. Under this condition, the inward gate of NqrD/E can flexibly open and close.

**Figure 6.**
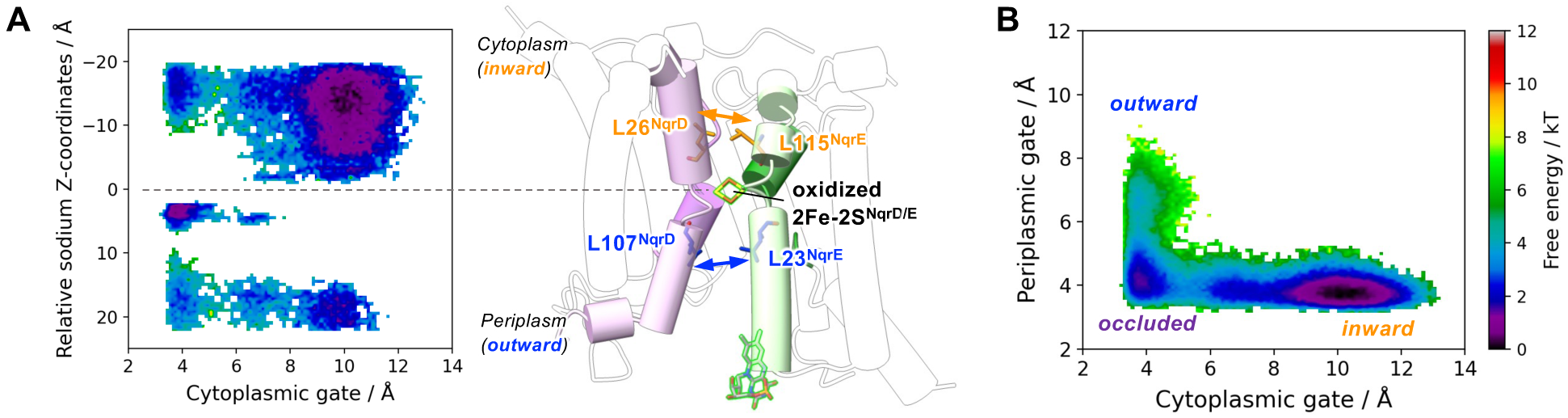
Free-energy analysis of NqrD/E conformational changes under the oxidized 2Fe-2S^NqrD/E^ condition. (A) Conformational free energy landscape projected on the axes of the cytoplasmic gate and relative z-coordinates of the Na^+^ being translocated in the transmembrane NqrD/E. (B) Conformational free energy landscape projected on the axes of the cytoplasmic and periplasmic gate sizes.

## Discussion

We have constructed redox-dependent MSMs to analyze the conformational free-energy landscape of the alternating-access mechanism of the NqrD/E subunits. The free-energy landscapes under different redox states of 2Fe–2S^NqrD/E^ and Na^+^-binding states suggest the following mechanism of conformational changes in NqrD/E during Na^+^-translocation (**Step1 - 3**, Figure 7).

**Figure 7.**
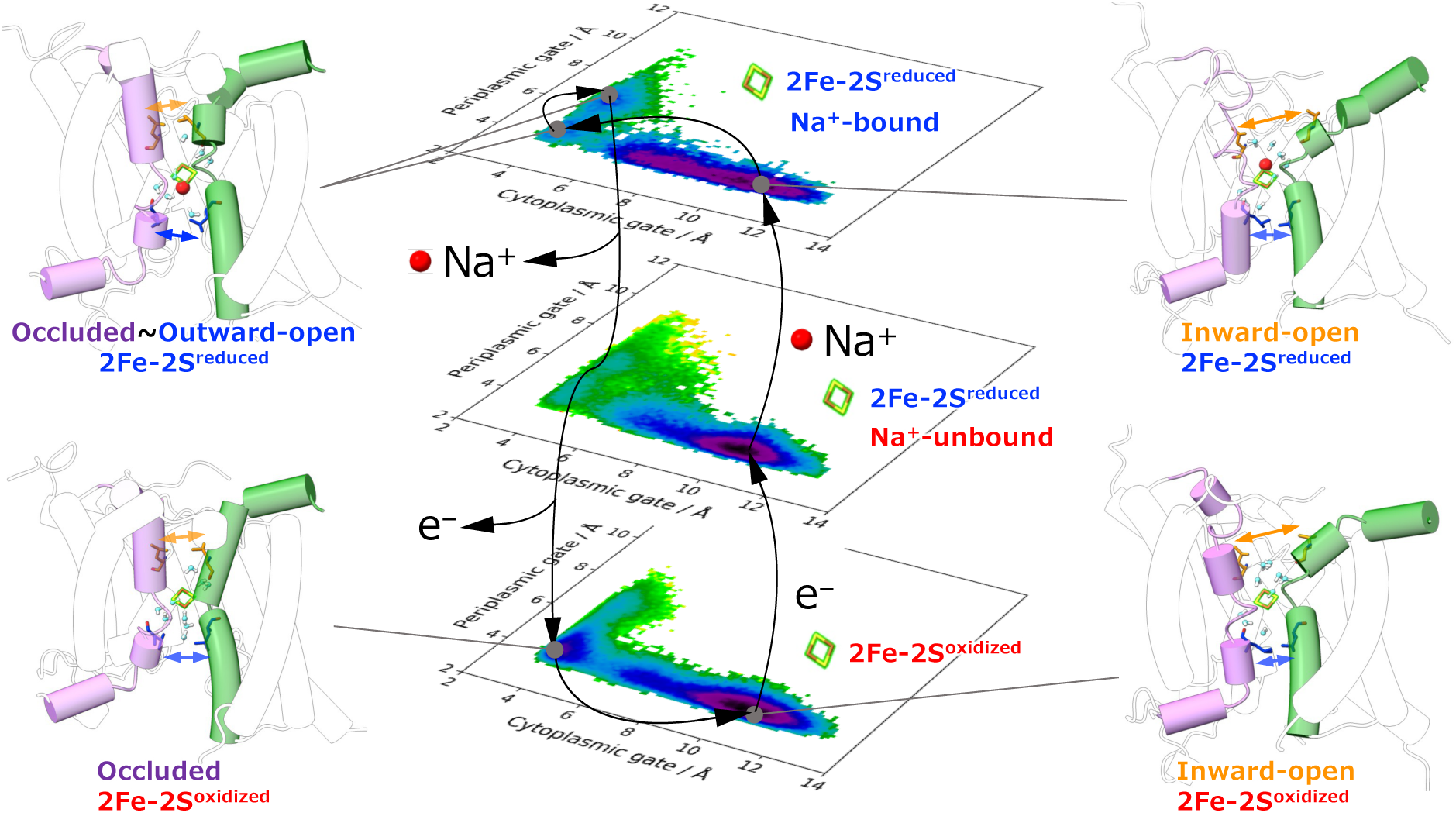
Proposed conformational cycles in NqrD/E of Na^+^-NQR during Na^+^ translocation driven by electron transfer.

**Step 1)** When 2Fe–2S^NqrD/E^ is in the oxidized state before the arrival of an electron, NqrD/E adopts either the inward-open or occluded conformation (Figure 7, bottom free energy landscape). After the arrival of an electron, the system transitions to the reduced 2Fe–2S^NqrD/E^ state (Figure 7, middle landscape).

**Step 2)** Upon reduction, 2Fe–2S^NqrD/E^ acquires a negative charge, destabilizing the occluded state and stabilizing the inward-open conformation. The negative charge also electrostatically attracts intracellular Na^+^. After Na^+^ binds near 2Fe–2S^NqrD/E^, the system transitions to the Na^+^-bound state (Figure 7, top landscape).

**Step 3)** In the reduced 2Fe–2S^NqrD/E^ and the Na^+^-bound state, NqrD/E can transition from the inward-open to the occluded state, and then to the outward-open state. The bound Na^+^ is released in the outward-open state. In the same conformational state, the transient approach of the next electron acceptor, FMN^NqrC^, on the periplasmic (extracellular) side toward 2Fe–2S^NqrD/E^ likely facilitates the subsequent inter-subunit electron transfer (Figure S3). The free energy landscape then returns to the bottom one shown in Figure 7, and the cycle restarts at **Step 1**.

Thus, the three energy landscapes derived from our MSM analyses collectively explain how redox changes and Na^+^-binding reshape NqrD/E’s conformational preferences to achieve directional Na^+^-pumping.

## Conclusions

In this study, we have developed an AlphaFold-facilitated Markov state modeling approach to investigating the redox-state-dependent Na^+^ translocation mechanism of Na^+^-NQR mediated by the NqrD/E alternating-access conformational change. The extensive MD-simulation data, initiated from diverse NqrD/E conformations generated by AlphaFold with MSA subsampling and with templates, and by adaptive sampling, enable the construction of redox-state-dependent MSMs. The MSMs allow quantitative analysis of the conformational free energy and transition pathway, revealing that the NqrD/E alternating-access conformational change is driven by Na^+^ binding near 2Fe–2S^NqrD/E^ in the reduced state and that this conformational change, rate-limited by the occluded-to-outward-open transition, carries Na^+^ across the membrane. This approach should prove useful for mechanistic investigations of electron- or proton-driven transporter proteins.

## Methods

### Simulation setup

All-atom MD simulations of Na^+^-NQR were performed using the cryo-EM structures (PDB IDs: 7XK3, 8A1U, 9UDF, and 9UD2), the crystal structure (PDB ID: 8ACW), and the AlphaFold structures as the initial structures. For the modeling of missing residues at the N-termini (for NqrB, NqrC, and NqrD) and C-termini (for NqrB and NqrC), the GalaxyFill plugin of CHARMM-GUI^37^ was used. The p*K*a of titratable residues was evaluated using PROPKA,^38^ and the protonation state was determined at pH 8, which is the same pH at which the experiments were conducted. The cofactors (FAD, 2Fe-2S^NqrF^, 2Fe-2S^NqrD/E^, FMN^NqrC^, FMN^NqrB^, and RBF^NqrB^) were retained as in the cryo-EM structures. FAD^NqrF^ is not covalently attached to NqrF. 2Fe-2S^NqrF^ is covalently attached to Cys70^NqrF^, Cys76^NqrF^, Cys79^NqrF^, and Cys111^NqrF^. The 2Fe-2S^NqrD/E^ is covalently attached to Cys29^NqrD^, Cys112^NqrD^, Cys26^NqrE^, and Cys120^NqrE^. FMN^NqrC^ is covalently attached to Thr225^NqrC^, while FMN^NqrB^ is attached to Thr236^NqrB^. RBF^NqrB^ is not covalently attached to NqrB.

The Na^+^-NQR structure was embedded in a 150 × 150 Å 1-Palmitoyl-2-oleoyl-sn-glycero-3-phosphocholine (POPC) membrane and solvated with TIP3P water and 150 mM sodium and chloride ions with the Membrane Builder plugin in CHARMM-GUI.^39,40^ The CHARMM36 force field^41^ was used except for the cofactors whose parameters were adapted from the works of Kaila group.^14,42,43^ The total number of atoms is ∼380,000.

### Molecular dynamics simulations

The systems were energy minimized with the restraint of non-hydrogen POPC, protein, and cofactor atoms for 10,000 steps. Then, they were equilibrated in three stages: (1) 100 ps equilibration at 150 K and constant volume with the restraint of non-hydrogen POPC, protein, and cofactor atoms, (2) 500 ps equilibration at 310 K and 1 bar pressure with the restraint of non-hydrogen protein and cofactor atoms, (3) 500 ps equilibration at 310 K and 1 bar pressure with the restraint of protein-Cα and non-hydrogen cofactor atoms. The restraint force constant was 1.0 kcal mol^-1^ Å^-2^ for each atom. The production runs were performed at 310 K and 1 bar pressure without restraints. Long-range electrostatic interactions were calculated by the particle mesh Ewald method^44^ with a direct space cut off of 12 Å. Langevin dynamics with 1 ps^-1^ damping coefficient was used for temperature control at 310 K, and the Nosé-Hoover Langevin piston was used for pressure control at 1 bar.^45–47^ The integration timestep was set to 2 fs by applying the SHAKE method^48^ to bonds involving hydrogen. All MD simulations were performed using NAMD 2.12 / 2.14 / 3.0.^49^ VMD,^50^ UCSF Chimera,^51^ UCSF ChimeraX,^52^ MDAnalysis,^53^ and MDTraj^54^ have been used for the analysis.

### Determination of the input features describing conformational changes in NqrD/E

First, MD snapshot structures were classified into inward-open or outward-open states as follows. If the minimum heavy-atom distance between residues Leu26^NqrD^ and Leu115^NqrE^ was > 4.5 Å and that between Leu107^NqrD^ and Leu23^NqrE^ was < 4.5 Å, they were classified into the inward-open state. Those with the opposite pattern (< 4.5 Å for Leu26^NqrD^ and Leu115^NqrE^ and > 4.5 Å for Leu107^NqrD^ and Leu23^NqrE^) were classified into the outward-open state. Then, residue-residue contacts were defined based on the minimum heavy-atom distance between residues *i* and *j* in each MD snapshot. If this distance was < 4.5 Å, the residues were considered to be in contact. The contact formation probability for each residue pair was calculated as the fraction of frames in which the contact was formed over the total number of frames. The contact formation probabilities scalculated within each conformational state are denoted as 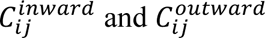, respectively. Residue pairs with 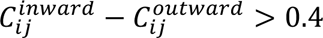 were classified as inward-open-specific contacts, whereas pairs with 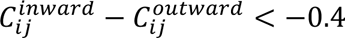 were classified as outward-open-specific contacts. The identified state-specific contacts were listed in Table S2. The minimum heavy-atom distances of the state-specific contacts are used as input features to describe conformational changes in NqrD/E.

### Markov state modeling

To construct a Markov state model from MD trajectories, the protein conformations were represented by 43 residue-pair distances as input features, derived from specific contacts found in the inward-open or the outward-open state, as shown in Table S2. These features were then subjected to tICA to reduce the dimensionality while preserving the slow collective motions of the alternating-access conformational changes in NqrD/E. Then, k-means clustering was applied in the tICA space to discretize the protein conformations into 100 discrete microstates. The number of microstates and the number of tICs were systematically evaluated using the VAMP-2 score,^31^ defined as the sum of the squared eigenvalues 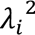 of the transition matrix whose elements correspond to the transition probabilities between states. Based on the convergence behavior of the VAMP-2 score, 100 microstates and 20 tICs were selected as optimal model parameters (Figure S1B). Implied timescales *t_i_*, which characterize the relaxation times of inter-state transitions, were then calculated from *λ_i_* as *t_i_* = − *τ*/*ln*|*λ_i_*|, and used to assess their dependence on the MSM lag time *τ*. The implied timescales converged for lag times of 4 ns or longer, indicating Markovian behavior (Figure S1C). Accordingly, a lag time of 4 ns was adopted for the final MSM. At this lag time, we also validated the reliability of the constructed MSM. The PCCA+ algorithm^32^ was applied to coarse-grain the 100 microstates to three macrostates (Figure S1E), and the resulting MSM was validated with the Chapman-Kolmogorov test, confirming that the transition probabilities were accurately estimated (Figure S1F). For transition path analysis, the MSM was coarse-grained into 31 macrostates using PCCA+. After defining the initial (inward-open state) and final (outward-open state) as groups of the macrostates, transition path theory^33^ was applied to compute the reactive flux *F_ij_* from microstate *i* to microstate *j*. The net-reactive flux was then calculated as 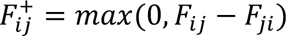. The transition pathways were subsequently identified by pathway decomposition based on 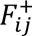. For each pathway *q*, its contribution was quantified by the minimum net-reactive flux along the pathway, **f*_q_*, which was used to compute the pathway probability *p_q_* = *f_q_*⁄∑*_r_ f_r_*.^33^ Relative net-reactive flux values on the primary transition pathway were computed as 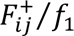 (Figure 5), where **f*_1_* represents the minimum net-reactive flux along the primary pathway. PyEMMA was used to construct and analyze MSMs.^55^

## Acknowledgements

The computations were performed using the Research Center for Computational Science, Okazaki, Japan (Projects 23-IMS-C201, 24-IMS-C198, and 25-IMS-C227). This work was supported by JSPS KAKENHI Grant Numbers JP23K14160 (to J.O.), JP22H02595, JP23K23858, and JP25H02299 (to K.O.).

