## Supporting information for "Conformational Dynamics of Na^+^-Pumping NADH-Quinone Oxidoreductase during Na^+^ Translocation from AlphaFold-Facilitated Markov State Modeling"

AUTHOR INFORMATION

**Corresponding Author**

*

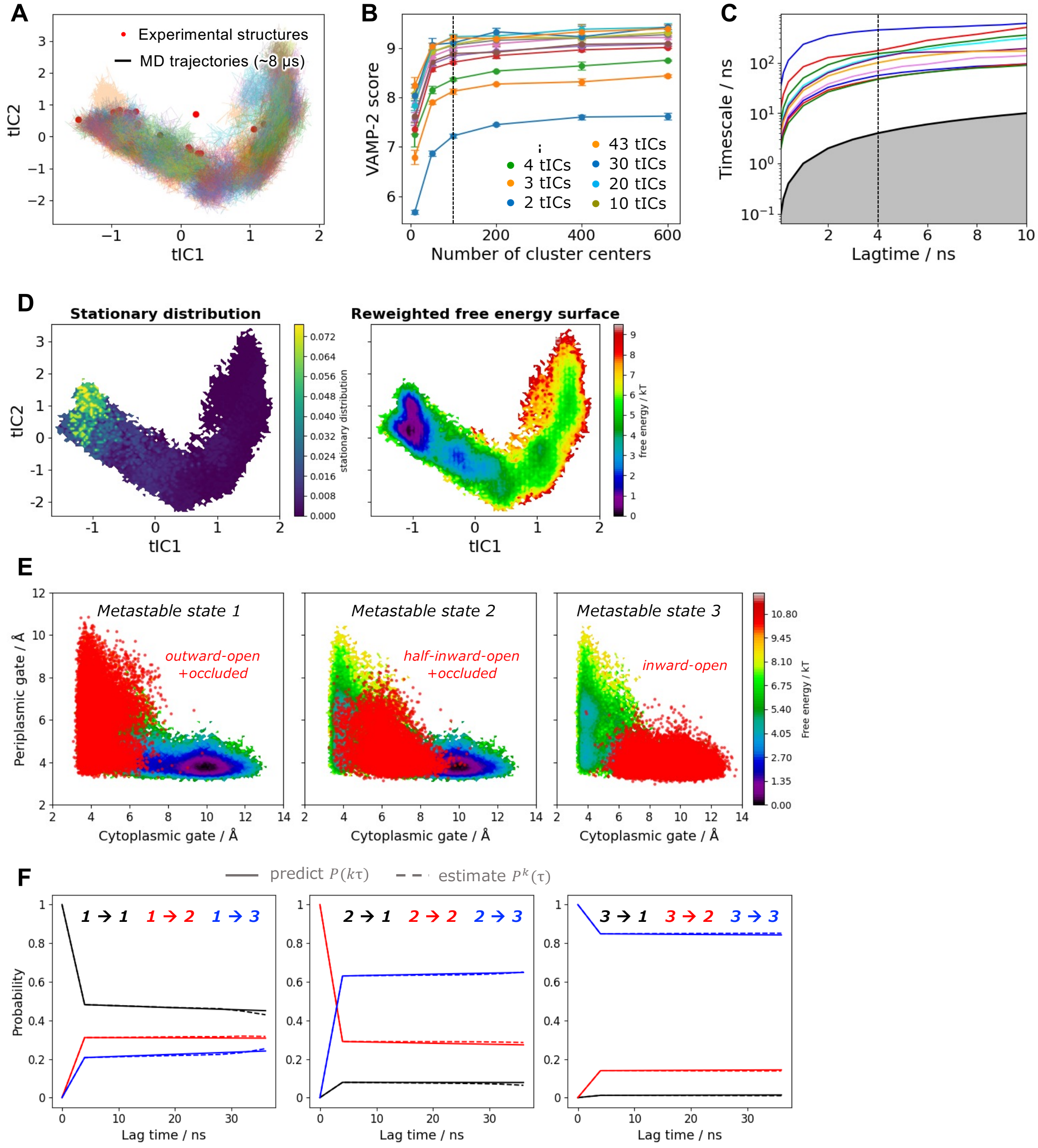


**Figure S1.** Construction and validation of the Markov state model (MSM) under the reduced 2Fe-2S$^{NqrD/E}$ condition. (A) MD trajectories used for Markov state modeling projected onto tICA coordinates. (B) Evaluation of our MSM using the VAMP-2 score. The model with 100 microstates and 20 time-lagged independent components (tICs) achieved a convergence of the score and was therefore selected for subsequent MSM analyses. (C) The implied timescale $t_i = -\tau/ln|\lambda_i|$ of MSM with the lag time $\tau$ and eigenvalue $\lambda_i$ of the transition matrix is plotted as a function of the lag time $\tau$. The implied timescales converged at the lag time of 4 ns, which was therefore used in the subsequent MSM analyses. (D) Equilibrium distribution and reweighted free energy landscape projected on the axis of tICs. (E) Distributions of three coarse-grain states computed with PCCA+ for a Chapman-Kolmogorov test. (F) Chapman-Kolmogorov test. The predicted probability $p(k\tau)$ and the estimated probability $p^k(\tau)$ are shown in the solid and dotted lines, respectively. Since these probabilities are almost identical, it was confirmed that our MSM correctly estimates the transition probabilities between states.

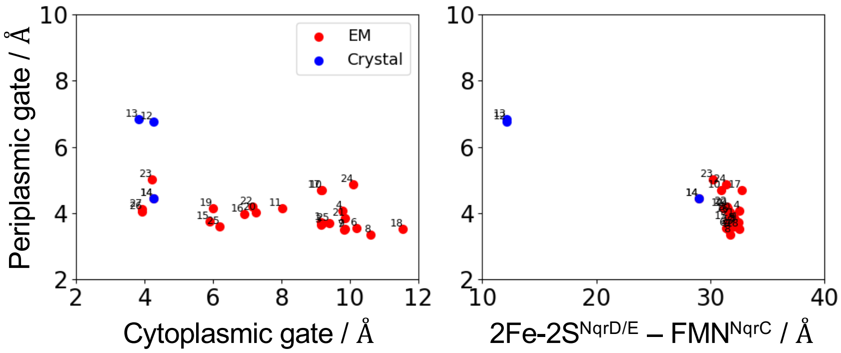


| Index | State: PDB ID |
|---|---|
| 1 | WT$_{ox}$ state 1: 7XK3 |
| 2 | WT$_{ox}$ state 2: 7XK4 |
| 3 | WT$_{ox}$ state 3: 7XK5 |
| 4 | WT$_{ox}$ + AD-42: 7XK6 |
| 5 | WT$_{ox}$ + KR: 7XK7 |
| 6 | WT$_{ox}$ : 8A1T |
| 7 | WT$_{red}$ + UQ2: 8A1U |
| 8 | WT$_{ox}$ + UQ2: 8A1V |
| 9 | WT$_{ox}$ + UQ1: 8A1W |
| 10 | WT$_{ox}$ + DQA: 8A1X |
| 11 | WT$_{ox}$ + HQNO: 8A1Y |
| 12 | WT$_{ox}$_1: 8ACW (crystal) |
| 13 | WT$_{ox}$_2: 8ACY (crystal) |
| 14 | WT$_{ox}$_3: 8AD0 (crystal) |

| Index | State: PDB ID |
|---|---|
| 15 | WT$_{red}$ : 9UUU |
| 16 | WT$_{red}$ w/o $Na^+$ (*up*): 9UD6 |
| 17 | WT$_{red}$ w/o $Na^+$ (*middle*): 9UD8 |
| 18 | WT$_{red}$ w/o $Na^+$ (*down*): 9UD9 |
| 19 | WT$_{red}$ + KR: 9UD5 |
| 20 | WT$_{red}$ + AD-42: 9UDG |
| 21 | G141A$^{NqrB}_{ox}$ + KR: 9LRR |
| 22 | G141A$^{NqrB}_{red}$ + KR (*stable*): 9UDA |
| 23 | G141A$^{NqrB}_{red}$ + KR (*shifted*): 9UDF |
| 24 | T236Y$^{NqrB}_{ox}$ : 9UD3 |
| 25 | T236Y$^{NqrB}_{red}$ : 9UD4 |
| 26 | T225Y$^{NqrC}_{ox}$ : 9U5G |
| 27 | T225Y$^{NqrC}_{red}$ : 9UD2 |

**Figure S2.** Experimental structures. The *upper-left* panel shows a 2D plot of the distances for the inward and outward gates. The inward and outward gate sizes are defined by the closest heavy atom distances: between Leu26$^{NqrD}$ and Leu115$^{NqrE}$ for the inward gate, and between Leu107$^{NqrD}$ and Leu23$^{NqrE}$ for the outward gate. The *upper-right* panel displays a 2D plot illustrating the closest heavy atom distance between the cofactors 2Fe-2S$^{NqrD/E}$ and FMN$^{NqrC}$, along with the outward gate distance.

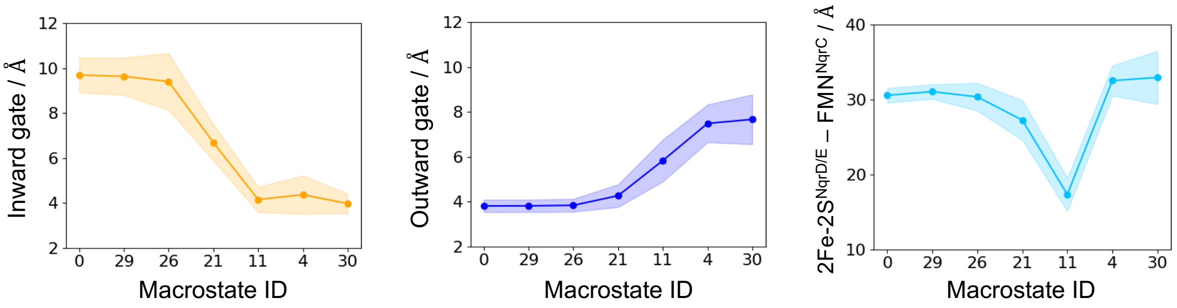


**Figure S3.** Progressions of inward (cytoplasmic), outward (periplasmic) gate sizes, and cofactor distance on the primary transition pathway from transition path analysis. The *left* and *center* panels show inward and outward gate sizes, respectively. The *right* panel shows the closest heavy-atom distance between 2Fe–2S$^{NqrD/E}$ and FMN$^{NqrC}$. In each panel, dots represent the mean, and the shaded area indicates the mean ± standard deviation.

| Initial structures | NqrD/E conformations | **Length** / ns | **Total length** / μs |
|---|---|---|---|
| Experiments | Inward-open<br>(PDB IDs: 7XK3 and 8A1U) | **1,000**<br>**1,000** | **2.6** |
| | Occluded<br>(PDB ID: 9UD2) | **400** | |
| | Half-outward-open<br>(PDB ID: 9UDF) | **200** | |
| AlphaFold2 | Half-outward-open | **400** | **1.2** |
| AlphaFold3 | Half-inward-open | **200** (40×5) | |
| | Half-inward-open | **200** (40×5) | |
| | Half-inward-open | **200** (40×5) | |
| | Occluded | **200** (40×5) | |

**Table S1. List of MD simulations (reduced 2Fe-2S$^{NqrD/E}$) initiated from experimental and AlphaFold structures for Markov state modeling.** MD trajectories initiated from AF3 structures were added to the dataset after the first adaptive-sampling cycle. For the AlphaFold2-predicted structures, which contain only protein subunits, cofactors were embedded by superposition onto the experimental structure (PDB ID: 8A1U). Specifically, FAD$^{NqrF}$ and RBF$^{NqrB}$ were embedded by aligning all Cα atoms of the corresponding subunits, whereas 2Fe–2S$^{NqrF}$ and 2Fe–2S$^{NqrD/E}$ were embedded by aligning the coordinating sulfur atoms (Cys70$^{NqrF}$, Cys76$^{NqrF}$, Cys79$^{NqrF}$, and Cys111$^{NqrF}$ for 2Fe–2S$^{NqrF}$; Cys29$^{NqrD}$ , Cys112$^{NqrD}$, Cys26$^{NqrE}$, and Cys120$^{NqrE}$ for 2Fe–2S$^{NqrD/E}$). FMN$^{NqrC}$ and FMN$^{NqrB}$ were embedded by aligning Ser225$^{NqrC}$ and Ser236$^{NqrB}$, respectively.

| Pair ID | Residue 1 | Residue 2 | Pair ID | Residue 1 | Residue 2 |
|---|---|---|---|---|---|
| 1[a] | Asp44$^{\text{NqrC}}$ | Tyr168$^{\text{NqrC}}$ | 25[b] | Ala22$^{\text{NqrD}}$ | Gln205$^{\text{NqrD}}$ |
| 2[a] | Ala33$^{\text{NqrD}}$ | Phe104$^{\text{NqrD}}$ | 26[b] | Leu26$^{\text{NqrD}}$ | Leu115$^{\text{NqrE}}$ |
| 3[a] | Thr40$^{\text{NqrD}}$ | Gln100$^{\text{NqrD}}$ | 27[b] | Val43$^{\text{NqrD}}$ | Phe104$^{\text{NqrD}}$ |
| 4[a] | Val43$^{\text{NqrD}}$ | Ile97$^{\text{NqrD}}$ | 28[b] | Asn68$^{\text{NqrD}}$ | Gln92$^{\text{NqrE}}$ |
| 5[a] | Met44$^{\text{NqrD}}$ | Leu101$^{\text{NqrD}}$ | 29[b] | Val70$^{\text{NqrD}}$ | Gln92$^{\text{NqrE}}$ |
| 6[a] | Val51$^{\text{NqrD}}$ | Val86$^{\text{NqrD}}$ | 30[b] | Arg71$^{\text{NqrD}}$ | Gln92$^{\text{NqrE}}$ |
| 7[a] | Ile66$^{\text{NqrD}}$ | Val70$^{\text{NqrD}}$ | 31[b] | Arg71$^{\text{NqrD}}$ | Glu95$^{\text{NqrE}}$ |
| 8[a] | Arg71$^{\text{NqrD}}$ | Met123$^{\text{NqrD}}$ | 32[b] | Ile72$^{\text{NqrD}}$ | Val91$^{\text{NqrE}}$ |
| 9[a] | Gly106$^{\text{NqrD}}$ | Phe123$^{\text{NqrE}}$ | 33[b] | Ile72$^{\text{NqrD}}$ | Pro114$^{\text{NqrE}}$ |
| 10[a] | Leu107$^{\text{NqrD}}$ | Leu23$^{\text{NqrE}}$ | 34[b] | Asn111$^{\text{NqrD}}$ | Val118$^{\text{NqrE}}$ |
| 11[a] | Thr110$^{\text{NqrD}}$ | Asn119$^{\text{NqrE}}$ | 35[b] | Cys112$^{\text{NqrD}}$ | Leu115$^{\text{NqrE}}$ |
| 12[a] | Thr110$^{\text{NqrD}}$ | Cys120$^{\text{NqrE}}$ | 36[b] | Cys112$^{\text{NqrD}}$ | Val118$^{\text{NqrE}}$ |
| 13[a] | Thr27$^{\text{NqrE}}$ | Leu115$^{\text{NqrE}}$ | 37[b] | Met115$^{\text{NqrD}}$ | Pro114$^{\text{NqrE}}$ |
| 14[a] | Val31$^{\text{NqrE}}$ | Leu115$^{\text{NqrE}}$ | 38[b] | Glu119$^{\text{NqrD}}$ | Pro114$^{\text{NqrE}}$ |
| 15[a] | Thr37$^{\text{NqrE}}$ | Phe112$^{\text{NqrE}}$ | 39[b] | Thr27$^{\text{NqrE}}$ | Phe112$^{\text{NqrE}}$ |
| 16[a] | Gly40$^{\text{NqrE}}$ | Leu109$^{\text{NqrE}}$ | 40[b] | Ala30$^{\text{NqrE}}$ | Phe112$^{\text{NqrE}}$ |
| 17[a] | Gly40$^{\text{NqrE}}$ | Phe112$^{\text{NqrE}}$ | 41[b] | Val31$^{\text{NqrE}}$ | Leu109$^{\text{NqrE}}$ |
| 18[a] | Leu41$^{\text{NqrE}}$ | Phe112$^{\text{NqrE}}$ | 42[b] | Thr37$^{\text{NqrE}}$ | Ala108$^{\text{NqrE}}$ |
| 19[a] | Ala44$^{\text{NqrE}}$ | Leu109$^{\text{NqrE}}$ | 43[b] | Leu41$^{\text{NqrE}}$ | Leu109$^{\text{NqrE}}$ |
| 20[a] | Ala44$^{\text{NqrE}}$ | Ile116$^{\text{NqrE}}$ | | | |
| 21[a] | Val45$^{\text{NqrE}}$ | Ile116$^{\text{NqrE}}$ | | | |
| 22[a] | Val48$^{\text{NqrE}}$ | Ile116$^{\text{NqrE}}$ | | | |
| 23[a] | Ile111$^{\text{NqrE}}$ | Leu91$^{\text{NqrF}}$ | | | |
| 24[a] | Met129$^{\text{NqrE}}$ | Asp133$^{\text{NqrE}}$ | | | |

**Table S2. List of state-specific contacts whose distances are used as input features describing conformational changes in NqrD/E.**

[a] Contact pairs specifically found in the *inward-open* state.

[b] Contact pairs specifically found in the *outward-open* state.

| Adaptive round | **Length** / ns | **Total length** / μs |
|---|---|---|
| 1 | **1,000** (**40**×25) | |
| 2 | **2,000** (**40**×50) | **4.0** |
| 3 | **1,000** (**40**×25) | |

**Table S3. List of MD simulations (reduced 2Fe-2S$^{NqrD/E}$) from adaptive sampling for Markov state modeling.**
